# PEX11β-mediated improvement of mitochondrial dysfunction restores behavioral defect and cellular viability in neuropathological conditions

**DOI:** 10.1101/2025.10.24.684183

**Authors:** Vaishali Kumar, Pradeep Kodam, Chaitanya Kunja, Pragya Komal, Amartya Sanyal, Shuvadeep Maity

## Abstract

Peroxisomal biogenesis factor 11 beta (Pex11β) is a key regulator of peroxisome proliferation, functioning in coordination with mitochondrial fission proteins. Recent human proteome co-regulation mapping has revealed a distinctive co-expression pattern between Pex11β and components of the mitochondrial respiratory chain complex at the peroxisome–mitochondria interface. In neurodegenerative disorders, such as Huntington’s disease (HD) and Alzheimer’s disease (AD), mitochondrial dysfunction is primarily associated with impairments in the respiratory chain complexes, particularly Complex II. However, the role of Pex11β in maintaining mitochondrial homeostasis in these two disease conditions remains largely unexplored. In this study, using both *in vitro* and *in vivo* models, we identify Pex11β as a critical modulator of mitochondrial dynamics, influencing the balance between fission and fusion. Through targeted knockdown and overexpression experiments, we demonstrate a direct link between Pex11β level and mitochondrial dysfunction, mediated by transcriptional dysregulation of nuclear-encoded Complex II and mitochondrial fusion genes. Restoration of Pex11β expression, along with Complex II and fusion gene level in striatal brain via a peroxisome proliferator & chemical chaperone, significantly alleviates motor and cognitive deficits in 3-NP-induced HD mouse model indicating better mitochondrial health in striatal brain. Furthermore, *in silico* analysis and ChIP-qPCR reveal Yin Yang 1 (YY1) is a critical transcription factor governing the co-expression of Pex11β and Complex II genes. This establishes a novel transcriptional axis that orchestrates inter-organellar communication. Collectively, our findings position Pex11β as a pivotal mediator of mitochondrial fission-fusion dynamics, linking mitochondrial morphology and oxidative phosphorylation (OXPHOS) to behavioural outcomes, and offering mechanistic insights relevant to amyloid-associated neurodegenerative diseases.

## 1. Introduction

The human brain consumes more energy than any other organ (*1*). Remarkably, neurons account for approximately 70%-80% of the energy consumed by the brain as neurons rely predominantly on aerobic oxidative phosphorylation via mitochondria to meet their energy demands (*2*) (*3*). The oxidative phosphorylation (OXPHOS) system is essential for sustaining high-energy requirements necessary for proper neuronal function. Consequently, dysregulation of the OXPHOS system has been frequently implicated in various neurodegenerative diseases. Notably, brain tissues from patients with Huntington’s disease (HD) and Alzheimer’s disease (AD) exhibit significant impairments in mitochondrial respiratory complexes, particularly complex II (*4*) (*5*) (*6*). These defects are accompanied by alterations in mitochondrial dynamics, membrane potential, and elevated production of reactive oxygen species (ROS), - hallmarks of mitochondrial dysfunction.

Compelling evidence indicates that mitochondrial dysfunction in HD and AD is driven by the accumulation of protein aggregates, including mutant huntingtin (mtHtt) and β-amyloid protein (*7*) (*8*) (*9*) (*10*). Notably, detailed analysis of genetic and pharmacological HD models has revealed deeper insights into the relationship between mitochondrial dysfunction and the resulting neurological and behavioral impairments. Post-mortem analysis of HD patients’ brains, along with various *in vivo* and *in vitro* HD models expressing either mutant full-length or N-terminal fragments of mutant Htt, consistently demonstrated morphological and functional abnormalities in mitochondria. These defects are closely associated with altered expression of mitochondrial fusion and fission proteins (*4*)(*5*). Like genetic mouse HD models, 3-nitropropionic acid (3-NP, a complex II inhibitor) induce a chemical HD model that recapitulates the key behavioral symptoms of HD with severe mitochondrial dysfunction, despite the absence of mutant Htt aggregates. The chemical HD model displays two hallmark features of HD: mitochondrial fragmentation and behavioral defects. All these findings suggest a shared mechanistic link across genetic and chemical HD models, wherein Complex II dysfunction and disrupted mitochondrial dynamics converge to drive disease pathology, regardless of the presence of protein aggregates. Similar observations regarding mitochondrial dysfunction in AD were reported previously (*10*). Mitochondria-shaping proteins regulate mitochondrial dynamics through cycles of fusion and fission, enabling the organelle to adapt its structure in response to cellular energy demands. Their dynamics are essential for the redistribution of healthy and damaged mitochondria, thereby maintaining cellular homeostasis (*11*) (*12*) (*13*). Depending on the cellular context—such as division, metabolic state, or pathological conditions—mitochondria may undergo fission to form smaller fragments or fusion to generate elongated networks. Disruption of mitochondrial fusion impairs mitochondrial function, including a loss of respiratory capacity, as observed in both yeast and mammalian cells (*14*). In mammals, mitochondrial fission is orchestrated by the cytosolic protein DRP1/DNM1L (dynamin-related protein 1), which is recruited to constriction sites by fission factors such as mitochondrial fission factor (MFF), mitochondrial fission 1 (FIS1). Conversely, mitochondrial fusion is mediated by mitofusins (MFN1, MFN2) located on the outer mitochondrial Membrane (OMM), and optic atrophy 1 (OPA1) located on the inner mitochondrial membrane (IMM) in mammalian cells.

Interestingly, peroxisome fission and proliferation also require the same mitochondrial fission machineries– DRP1, MFF, FIS1–alongside the peroxisomal protein, PEX11, particularly the β isoform (PEX11β), highlighting a shared regulatory axis between these two organelles. Recent studies have shown that overexpression of MFNs not only induces mitochondrial clustering but also promotes co-clustering of peroxisomes (*15*). This observation highlights a complex interdependence between these organelles, mediated by mitochondrial fission-fusion proteins, with PEX11β emerging as a potentially crucial player (*16*). A clinical study involving a patient with PEX11β mutation (c.743_744delTCinsA mutation in the exon 4 of the PEX11 β gene) associated with a subgroup of peroxisomal biogenesis disorders revealed neurological anomalies and abnormal gait, symptoms that overlap with behavioral features of HD (*17*). Remarkably, therapeutic intervention targeting mitochondrial function alleviated these symptoms, suggesting a functional interplay between PEX11β and mitochondrial dysfunction (*18*). Further supporting this connection, human proteome correlation mapping by Kustatscher et al. identified a unique co-expression regulation between OXPHOS proteins and PEX11β, but not with other PEX proteins (*19*). Despite these insights, the molecular role of PEX11β in modulating mitochondrial dysfunction in neurodegenerative diseases such as HD and AD remains largely unexplored.

In this study, we systematically identified the downregulation of PEX11β across post-mortem expression data sets from HD and AD patients. We further validated the co-expression pattern between PEX11β and complex II in both genetic and chemical models of HD. Using *in vitro* HD and AD models, we confirmed the involvement of PEX11β in controlling mitochondrial fragmentation. We also demonstrated that the chemical chaperone-like 4-PBA can improve motor and cognitive behavior in the 3-NP HD mouse model via restoration of PEX11β. Overall, our results established a direct and previously unreported involvement of PEX11β in HD and AD pathology. Furthermore, our study identified YY1-PEX11β-Complex II transcriptional axis regulating mitochondrial dynamics and mitochondrial-peroxisomal crosstalk.

In conclusion, consistent expression patterns of mitochondrial fission-fusion genes across HD and AD models suggest a shared molecular mechanism underlying these neurodegenerative conditions. Notably, restoring mitochondrial dynamics and enhancing cellular viability through the upregulation of PEX11β presents a promising therapeutic avenue for treating both disorders.

## 2. Results

### 2.1 Association of PEX11β with organellar fission in HD and AD conditions

HD is marked by the accumulation of mutant HTT (Huntingtin) aggregates in the cytoplasm of brain cells, while AD is characterized by intra-neuronal deposits of amyloid-beta (Aβ) aggregates, including oligomers, protofibrils (PFs), and mature fibrils. A common hallmark of both conditions is mitochondrial dysfunction accompanied by altered dynamics (*20*) (*21*).

To better understand the mechanisms underlying mitochondrial dysfunction, we conducted a comparative analysis using the MitoCarta3.0 database (comprising 1,136 mitochondrial genes) and differentially expressed genes (DEGs) identified from post-mortem brain transcriptomes of HD (1,082 genes) and AD (3,293 genes) (Figure 1A, Supplementary Figure 1A, Supplementary file III, see methods for details). Through this meta-analysis, we identified 209 differentially expressed DEGs directly associated with mitochondrial regulation, which we termed mitochondria-specific DEGs (mtDEGs). Pathway enrichment analysis using GO biological process revealed significant dysregulation in OXPHOS/respiratory complex (GO: 0045333, GO:0019646, GO:004275, GO:0009060, GO:0006119, GO:0006120)– ranking among the top 10 enriched pathways (Figure 1B, Supplementary File II). Notably, the expression of OXPHOS genes is markedly downregulated (Supplementary Figure 1C). Interestingly, among the 209 mtDEGs, the maximum proportion of genes (40%) are those that encodes for mitochondrial inner membrane proteins (Figure 1C). These findings are consistent with previous studies on OXPHOS gene dysregulation in HD and AD (*22*) (*23*) (*24*).

**Fig. 1.**
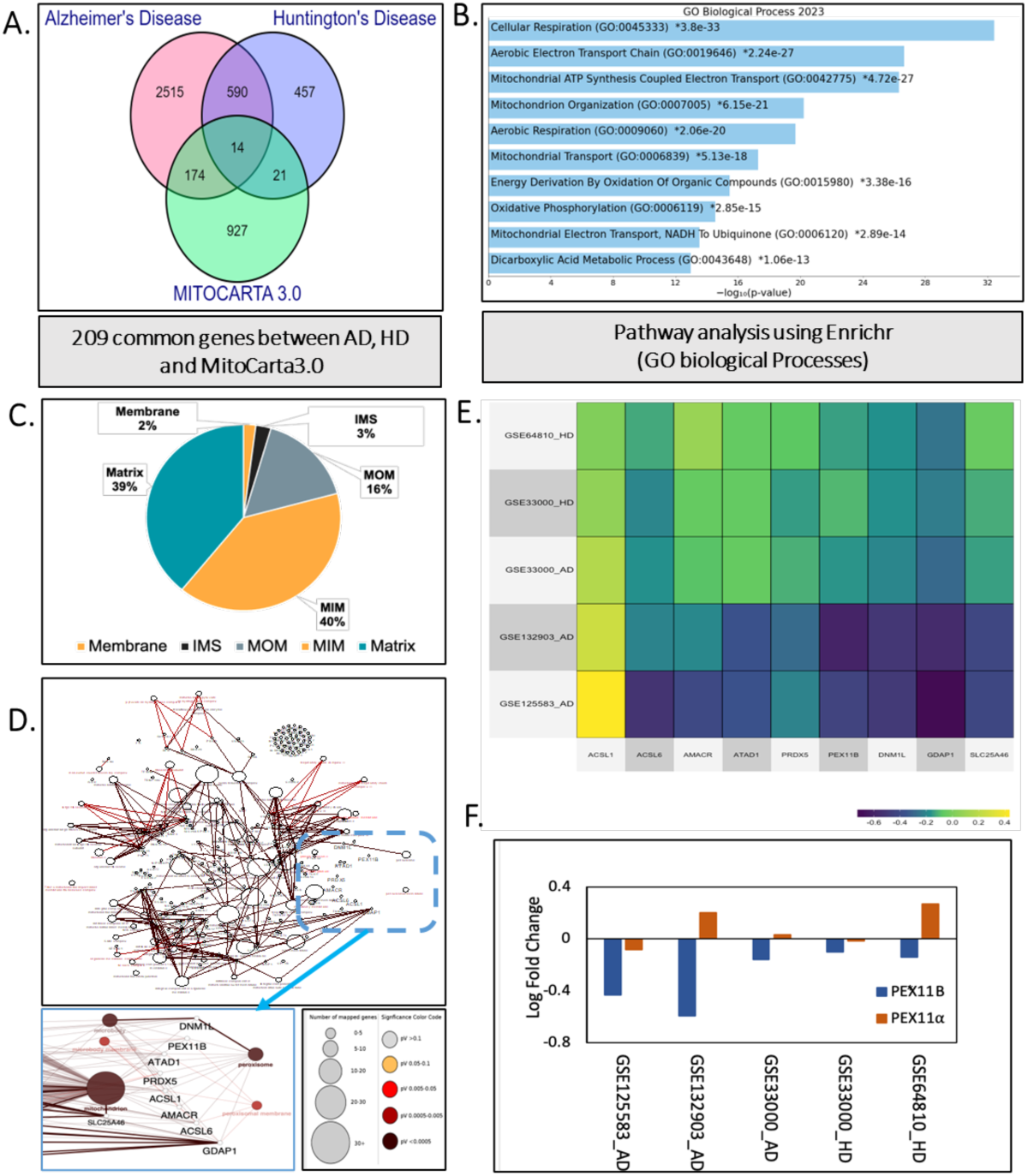
Decreased expression of PEX110 is associated with proteins controlling mitochondrial organisation in HD and AD conditions. (A) Venn overlap analysis among AD, HD, and MitoCarta3.0 database identified of 209 common genes shared between MitoCarta3.0 and AD or HD datasets. (B) Pathway Enrichment analysis of 209 genes using the Enrichr tool showed the top 10 significantly changed GO biological processes. (C) Systematic segregation of 209 genes based on their spatial distribution in mitochondria. (D) A mitochondrial protein network diagram of 209 common genes from AD, HD, and MitoCarta3.0 was created using Cytoscape as indicated in the methods. The network identified inter-organellar interaction displaying PEXlip and interacting proteins. The size of nodes indicates the number of mapped genes, where the color map corresponds to the statistically significant interactions. (E) The heat map showed the normalized expression levels for RNA (Log2Fold change) of selected genes known to interact with Pexlip and found in the protein interaction map in HD/AD patients compared to controls. More blue signifies enhanced downregulation, whereas yellow indicates upregulation. (F) Bar graphs represented the expression (Log2Fold change) trends of PEXlla/P across AD and HD datasets. PEX11B expression showed a significant decrease (except the marked one) in patients compared to healthy individuals.

Careful analysis of the GO Biological Process ontologies revealed significant changes in pathways related to mitochondrial organization (GO:0007005) and organellar fission-fusion processes (Organelle fission GO:0048285; Mitochondrial Fission GO:0000266; Peroxisome Fission GO:0016559) (Supplementary Figure 1B, Supplementary Table III). Genes associated with these categories play a crucial role in regulating mitochondrial dynamics through the control of their fission-fusion cycles. Notably, the peroxisomal fission protein, Peroxisome biogenesis factor 11β (PEX11β), was found to be enriched alongside key mitochondrial proteins such as OPA1, DNM1L, GDAP1, and SLC25A46 (Supplementary table IV B). Previous studies have shown that OPA1, DNM1L, GDAP1, and SLC25A46 are involved in mitochondrial dynamics, while PEX11 and GDAP1 are known to contribute to peroxisomal fission (*25*) (*26*) (*27*). To further investigate the role of PEX11β, we analyzed the PPI network. The resulting interaction map identified several key partners– GDAP1,

ATAD1, DRP1/DNM1L, PRDX5, ACSL1, ACSL6, and AMACR–distributed across organelles such as the peroxisome, microbody, and mitochondria (Figure 1D). Transcriptomic data of both HD and AD models revealed a general downregulation of genes encoding these interacting proteins, with the exception of ACSL1 (Figure 1E, Supplementary Table IV B). Previous studies have linked the dysregulation of these proteins to mitochondrial fragmentation, highlighting their role in mitochondrial dynamics (*28*) (*25*) (*27*) (*29*) (*30*) (*31*). Two independent studies have demonstrated that PEX11β directly contributes to mitochondria-peroxisome interactions by facilitating physical contact between the organelles, thereby maintaining their spatial clustering (*16*) (*19*). Further curation using the MitoCarta3.0 database confirmed the enrichment of PEX11β across various brain regions, supporting its association with mitochondrial proteins (Supplementary file III). We also examined the expression profiles of the two major PEX11 isoforms (α and β) across HD and AD transcriptomic datasets, which showed significant downregulation in both conditions (Figure 1F, Supplementary Figure 1D). Together, the *in silico* findings suggest a critical link between reduced PEX11β expression and the pathophysiology of HD/AD. To understand the underlying mechanisms, we conducted a systematic investigation of the PEX11β-mitochondria relationship in multiple HD models, followed by targeted experiments to validate the phenomenon in AD conditions.

### 2.2 *in vitro* HD condition recapitulates the expression defect of PEX11β and altered mitochondrial dynamics identified in *in silic*o analysis

PEX11β is an essential protein required for peroxisome fission. Interestingly, its function is supported by three additional proteins–FIS1, MFF, and DRP1/DLP1/DNM1L – which are also key regulators of mitochondrial fission (*16*). Conversely, the mitochondrial fusion protein MFN1/2 was found to increase the peroxisome-mitochondrial cluster (Huo et al, 2022). Therefore, we interrogated the expression patterns of PEX11β along with the mitochondrial fission-fusion proteins (FIS1, MFF, DRP1/DNM1L, MFN1/2, OPA1). To understand the underlying mechanisms, we conducted a systematic investigation of the PEX11β-mitochondria relationship in multiple HD models, followed by targeted experiments to validate the phenomenon in the AD conditions.

We first assessed their expression in *in vitro* HD models using two different cell lines: HEK293 cells stably expressing Q23/Q74 and N2A cells transiently expressing Q23/Q100 (see Methods for details) (Figure 2a). RT-qPCR analysis revealed a consistent downregulation of PEX11β in both disease models (Figure 2b), corroborating the *in silico* transcriptomic findings (Figure 1F). These results suggest that reduced expression of PEX11β may serve as a potential molecular marker for HD pathology.

**Figure 2:**
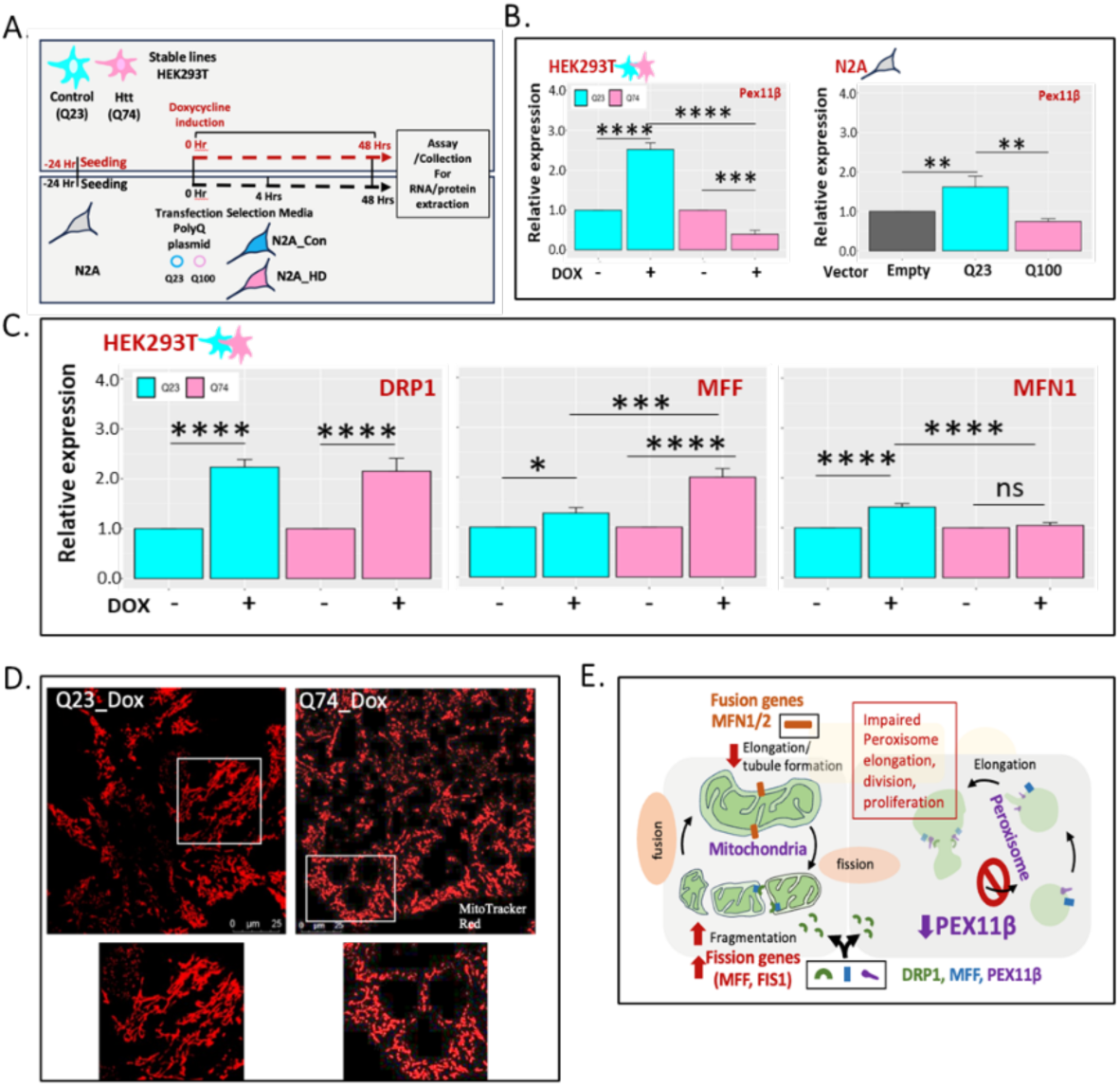
in vitro HD condition recapitulates the expression defect of PEX110 and altered mitochondrial dynamics. (A) Schematic representation of experimental outline showing dox-inducible stable lines (HEK293T) expressing either Poly-Q repeats (Q23/Control) or Poly-Q repeats (Q74/HD) huntingtin (HTT) exon 1 or N2A with transiently expressing Poly-Q repeats (Q23/Control & Q100/HD). Cells were collected for downstream assays, gene or protein expression studies, after the indicated times. (B) RT-qPCR confirms the decreased expression pattern of PEXllp in the HD condition, corroborating in silico transcriptomics data. (C) Fission genes (DRP1 and MFF) were expressed significantly higher in the HD condition, whereas no significant change in fusion gene (MFN1) expression was observed in the HD condition, indicating altered mitochondrial dynamics with enhanced fission. (D) Confocal images of mitochondria stained with MitroTracker Red showed severe mitochondrial fragmentation in the HD in vitro model expressing higher Poly-Q repeats (Q74) compared to its control harboring PolyQ repeat 23 (Q23). Imaging corroborated the qRT-PCR pattern of fission/fusion genes. (E) Diagrammatic representation showing altered expression pattern of fission fusion genes along with PEXllp, leading to impaired organellar (mitochondrial and peroxisomal) fission. Bar plot is from three independent experiments (n = 3) and all values are presented as mean with SD; *p value calculated with paired t-test with Holm’s correction *P < 0.05. *p < 0.05, **p < 0.005, ***p < 0.0005, and ****p < 0.00005).

Next, we examined the expression patterns of key mitochondrial fission and fusion regulators— MFF, DRP1/DNM1L, and MFN1/2—across HD models (Figure 2C, Supplementary Figure 3D). Among these, the fusion protein MFN1 was notably downregulated, while the fission protein MFF was significantly upregulated under HD pathological conditions. Although DRP1/DNM1L expression increased after 48 hours in both control (Q23-induced) and HD (Q74-induced) cells, MFF expression was markedly elevated only in the HD condition (Figure 3F). The expression pattern of fission and fusion genes in N2A expressing Q23/Q100 also followed a similar trend observed in HEK293 stably expressing Q23/Q74 (Supplementary figure 3C).

**Figure 3:**
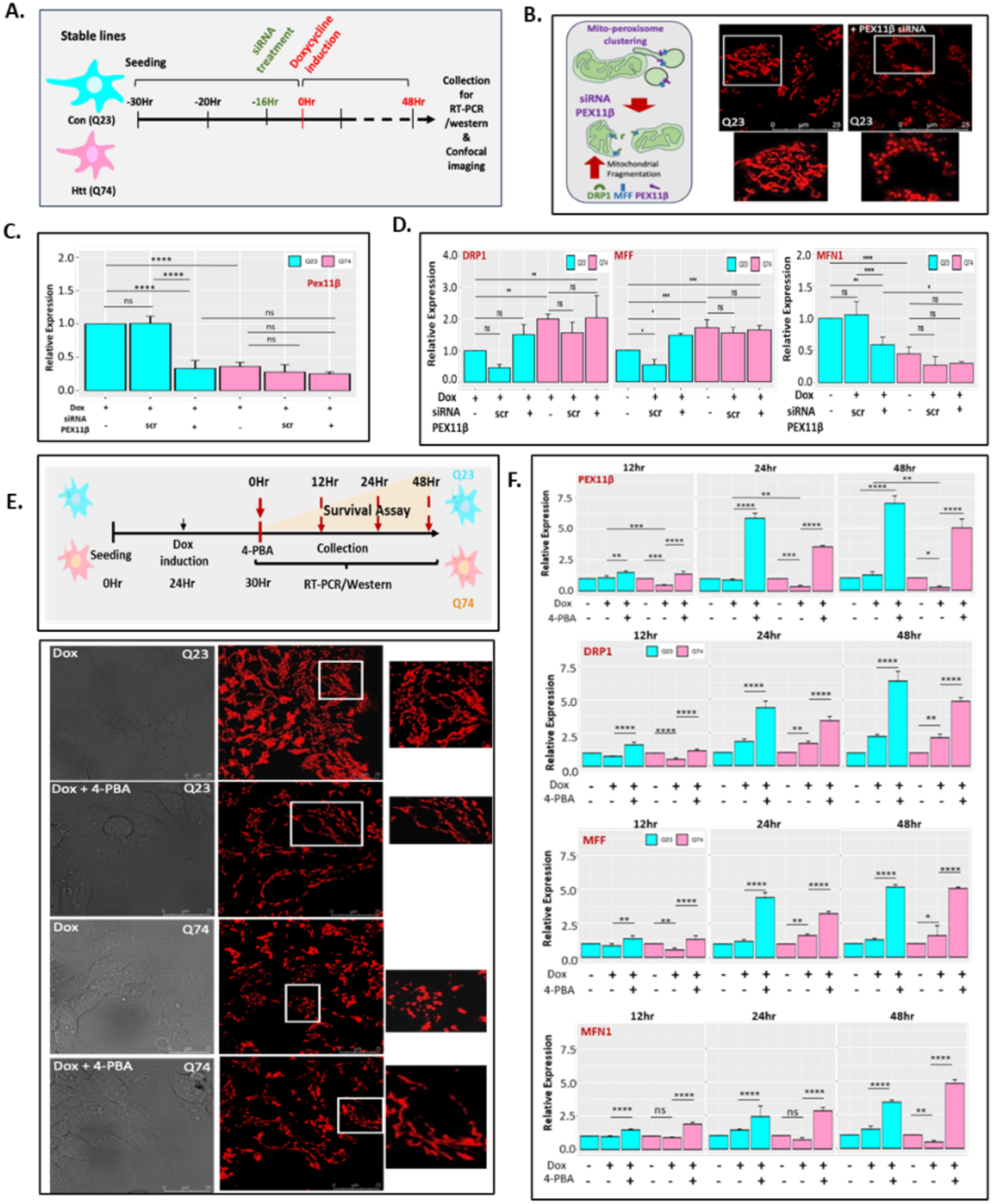
A**l**tered **mitochondrial dynamics under the control of PEXllf** (A) Schematic representation of the experimental outline showing siRNA-mediated for Pexllf knockdown to test its downstream effect. (B) Confocal imaging of MitoTracker-stained mitochondria showed enhanced fragmentation in Control cells after siRNA-mediated PEXllp knockdown. Schematic diagram showing mitochondrial fragmentation upon siRNA knockdown. (C) qRT-PCR confirmed reduced expression of PexliP after siRNA-mediated Pexllf knockdown in both Q23 and Q74 cells (n=3). (D) Gene expression study in Pexllf knockdown cells showed an increase in expression (MFF, DRP1), while the fusion gene showed a decreased expression (MFN1) in line with our imaging studies (n=3). (E) Schematic representation of the experimental outline of survival assay, with confocal imaging and RNA or protein collection after 4-PBA treatment at indicated time points (12, 24, 48 hours). Confocal imaging of MitoTracker-stained mitochondria showed rescue of mitochondrial morphology (more tubular) in Q74(HD) when treated with 4-PBA at 48 hours. (F) Bar plots across different time points after 4-PBA treatment showed significant enhancement of PEX11P expression Bar graph represents the unique expression pattern for genes (DNM1I/DRP1, MFN1, MFF) involved in mitochondrial dynamics after 4-PBA treatment. 4-PBA enhanced the expression of fission genes (DRP1 and MFF) and rescued the expression of the fusion gene MFN1 in Q74 (HD) (n=3). All gene expression data represent the mean (n = 3, biological replicates) with SD. *p-value calculated via paired t-test with “Bonferroni” correction.

Given that MFF is essential for recruiting DRP1 to mitochondria in mammalian cells, this selective upregulation likely enhances DRP1–MFF-mediated mitochondrial fission, contributing to the observed mitochondrial fragmentation in HD models (*33*) (*34*) (Itoyama et al., 2013). Additionally, previous studies have shown that mutant huntingtin (mtHtt) interacts with DRP1, further increasing its activity in HD (*36*).

Together with our in-silico findings, these in vitro expression studies support the notion of elevated mitochondrial fission in HD, with MFF emerging as a key contributor. Confocal imaging using MitoTracker Red further confirmed alterations in mitochondrial dynamics, revealing pronounced mitochondrial fragmentation in HD (Q74) cells compared to controls (Q23), consistent with the expression patterns of fission-related genes (Figure 2D, supplementary Figure 2A). These results collectively demonstrate that downregulation of PEX11β and upregulation of MFF are closely associated with HD pathology (Figure 2E).

### 2.3. Deletion of PEX11β impairs the fission-fusion process, resulting in fragmented mitochondria while overexpression of PEX11β restores a healthy mitochondrial network and cellular viability in HD conditions

Previous yeast genetic screening identified PEX11 as a critical component of the ERMES (Endoplasmic Reticulum-Mitochondria Encounter Structure) contact site, essential for establishing peroxisome– mitochondria interactions through the mitochondrial tethering protein MDM34 (*37*). Although direct evidence of PEX11β interacting with mitochondrial tethering proteins in mammals is currently lacking, a recent study demonstrated that overexpression of the mitochondrial fusion genes MFN1 and MFN2 leads to elongated mitochondria and increased peroxisome–mitochondria clustering (Huo et al, 2022).

To investigate whether PEX11β expression is linked directly to mitochondrial fission-fusion dynamics, we performed siRNA-mediated knockdown (KD) in control cells expressing a non-pathogenic polyQ repeat length (Q23). Confocal imaging revealed increased mitochondrial fragmentation in PEX11β-KD cells, indicating that PEX11β plays a direct role in maintaining elongated, fused mitochondrial structures (Figure 3A, B). Image quantification through MiNA showed an altered mitochondrial footprint in PEX11β-KD control cells indicating severe mitochondrial fragmentation (Supplementary Figure 3A). RT-qPCR analysis confirmed that a 50% reduction in PEX11β expression was sufficient to induce enhanced mitochondrial fragmentation in Q23 cells (Figure 3C).

Further, RT-qPCR of fission-related genes DRP1 and MFF supported the imaging data, showing elevated expression in PEX11β-KD Q23 cells (Figure 3D). In HD models, which already exhibit reduced PEX11β expression, fission gene expression was similarly elevated regardless of siRNA treatment. Notably, there was no significant difference in fission gene expression between control and HD cells under PEX11β-KD conditions, suggesting that knockdown of PEX11β does not further amplify fission gene expression in HD (Figure 3D). When we monitored the fusion gene MFN1 expression in PEX11β-KD cells, we found a consistent downregulation in both control and HD conditions. Importantly, the HD condition showed a more pronounced decrease in MFN1 expression compared to controls. The absence of PEX11β led to decreased expression of MFN1 in control cells which remained lower in the HD condition (Figure 3D). These findings suggest PEX11β is essential in maintaining fusion gene expression (particularly MFN1/2), not the fission genes, to preserve balanced mitochondrial dynamics. MFN1/2 deficiency, whether due to pathological conditions or experimental knockdown, disrupts this balance due to modulation of fusion genes.

If PEX11β is a key regulator of mitochondrial dynamics with concomitant expression of fusion genes, then enhancing its expression should also help in restoring fragmented mitochondrial morphology toward a more elongated network. To test this, we used 4-phenylbutyrate (4-PBA)—an FDA-approved chemical chaperone known to act as a peroxisome proliferator in a PPAR-independent manner—to upregulate endogenous PEX11β expression. Given that PEX11β interacts with mitochondrial fission proteins (MFF, DRP1/DNM1L, FIS1)—many of which exhibit dual localization—we assessed mitochondrial dynamics in HD cells with and without 4-PBA treatment. Confocal imaging using MitoTracker Red exhibited the restoration of elongated mitochondrial morphology after treatment with 4-PBA for 48 hours in HD (Q74-expressing) cells which usually harbor severely fragmented mitochondria (Figure 3E, Supplementary Figure 3E). We performed quantitative RT-PCR to confirm molecular-level changes after 4-PBA treatment. 4-PBA led to enhanced PEX11β expression over time (12, 24, and 48 hours) (Figure 3F). Interestingly, in HD cells, the expression of fission genes MFF and DRP1/DNM1L increased over time following 4-PBA exposure. However, the expression of the fusion gene MFN1, which was previously downregulated in HD, showed a marked increase after 4-PBA treatment (Figure 3F). Notably, the fold change in MFN1 expression exceeded that of the fission genes, suggesting a shift in mitochondrial dynamics toward a more fused, elongated phenotype. These findings were further validated in N2A cells transiently expressing Q23/Q100, which recapitulated the same expression patterns, confirming the conserved role of PEX11β in neuronal cells (Supplementary Figure 3C).

4-PBA has been previously used as a chemical chaperone to alleviate ER stress, a condition known for inducing mitochondrial fragmentation (*38*). Moreover, ER stress is widely recognized as a major pathological feature in HD. To determine whether the restoration of mitochondrial dynamics after 4-PBA treatment is a direct consequence of PEX11β transcriptional activation—rather than being a secondary effect of reduced ER stress—we examined mitochondrial morphology in PEX11β-KD cells treated with 4-PBA.

Confocal imaging revealed that 4-PBA treatment failed to rescue elongated mitochondrial morphology control cells lacking PEX11β, confirming that PEX11β is essential for maintaining mitochondrial structure (Supplementary Figure 3A). Consistently, qRT-PCR analysis showed that the expression of MFN1 was not restored in 4-PBA-treated HD cells lacking PEX11β (Supplementary Figure 3B).

To our knowledge, this is the first experimental evidence demonstrating that PEX11β is indispensable for maintaining healthy mitochondrial dynamics in HD. Both knockdown and disease-associated downregulation of PEX11β result in severe mitochondrial fragmentation.

### 2.4 A PEX11β-dependent co-expression dysregulation between complex II and PEX11β is a unique transcriptional nexus in HD

Observing the unique transcriptional correlation of PEX11β with fusion genes controlling mitochondrial dynamics prompted us to investigate how mitochondrial genes could be linked with PEX11β, the key player in peroxisomal division and proliferation. Earlier studies showed that mitochondrial OXPHOS deficiencies, have been shown to impair peroxisomal proliferation through transcriptional remodeling mechanisms, highlighting a functional interdependence between these organelles (*39*). Supporting this same notion, a recent human proteome co-regulation study identified a unique co-expression pattern between OXPHOS components and PEX11β, suggesting a coordinated regulatory relationship (*19*) (*40*).

To further investigate this link between PEX11β, mitochondrial dysfunction, and OXPHOS dysregulation, we checked the expression profiles of genes encoding the five mitochondrial OXPHOS complexes across HD and AD datasets. Our *in silico* analysis revealed a consistent downregulation of these genes in both disease conditions (Supplementary Figure 1C), indicating significant disruption of OXPHOS function. These findings are consistent with earlier studies that reported mitochondrial respiratory chain impairments as a hallmark of both HD and AD pathologies (*22*) (*23*) (*24*).

Among these, Complex II dysregulation is a well-documented hallmark of HD, where the degeneration of medium spiny motor neurons has been directly linked to complex II inhibition. Targeted silencing of SDHB, a key complex II subunit, has been shown to induce mitochondrial dysfunction through elevated ROS production, ultimately leading to neuronal cell death (*41*). We thus focused on first evaluating both transcript and protein expression levels of complex II subunits—SDHA and SDHB at different time points in the HD cellular model expressing 74 polyglutamine (Q74) repeats. Our *in vitro* study corroborates the *in silico* transcriptomic data (Figure 4A, Supplementary Figure 4A-C). Notably, this reduction in complex II expression was correlated with significant downregulation of PEX11β at both the transcript and protein levels in the same HD model (Supplementary Figure 4A). The same expression pattern was observed in N2A transiently expressing toxic PolyQ repeats (Q100) (Figure 4A). To ensure complex II inhibition is dependent on PEX11β, we monitored expression of SDHB in the absence of PEX11β. PEX11β-KD cells showed decreased expression of SDHB which failed to restore even in the presence of 4-PBA (Figure 4B). These findings support the existence of a critical co-expression axis between PEX11β and complex II, wherein HD-associated downregulation of both components is linked with mitochondrial dynamics.

**Figure 4:**
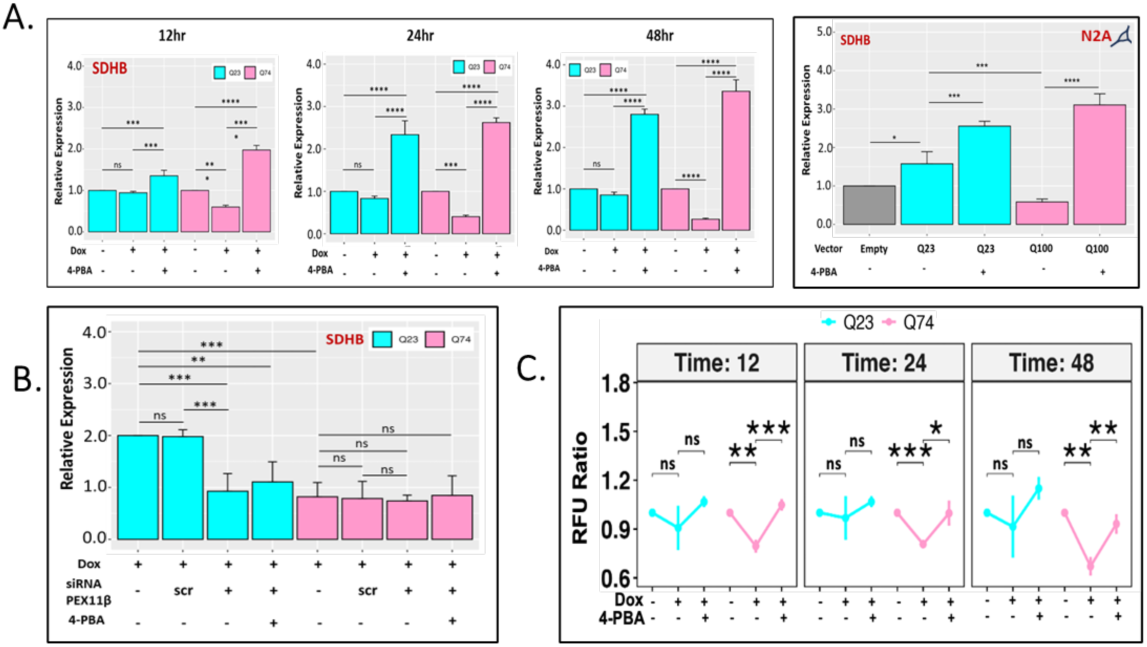
R**e**duced **expression of SDHB (Complex II subunit) in the HD condition was restored by 4-PBA treatment** (A) qRT-PCR confirmed reduced expression of SDHB in Q74 (HD) cells compared to Q23 (control) cells across three (12, 24, and 48 hours) different time points, while the expression of SDHB increased after 4-PBA treatment (n=3). Neuro2A cell line harbouring PolyQ repeats (Q23 or 0.100) displayed reduced SDHB expression in 0100 (HD) cells. (B) Q23/Q74 after siRNA-mediated Pexllp knockdown exhibited low expression of SDHB both in the presence or absence of 4-PBA. All gene expression data from A and B represented mean (n = 3, biological replicates) with SD. *p value calculated via paired t-test with “Bonferroni” correction) (C) Alamar Blue cell viability assay demonstrated recuse of cellular viability in Q74 (HD) cells after 4-PBA treatment at indicated times. *p value calculated via paired t-testJ* represents following p values: *p < 0.05, **p < 0.005, ***p < 0.0005, and ****p < 0.00005). Also, see supplementary figure 4 for expression data of other genes (SDHA, Mirol).

As a peroxisomal protein, PEX11β plays a critical role in maintaining peroxisomal dynamics, cooperating in fatty acid β-oxidation, ROS homeostasis, supporting local energy turnover and regulating mitochondria-dependent apoptosis (*40*) (*19*). Miro1, a membrane adaptor for the microtubule-dependent motors kinesin and dynein 60, known to be co-regulated with PEX11β found to be distributed both in mitochondria and peroxisomes (*19*) (*42*) (*43*). We confirmed a similar co-expression pattern of Miro1(supplementary Figure 4C). Thus, in HD condition, decreased expression of Complex II, PEX11β, and Miro1 resulted in severe mitochondrial dysfunction with altered dynamics, enhanced ROS production, altered peroxisome - mitochondrial clustering, followed by cellular death.

Building on this, we investigated whether the cell death observed in HD could be linked to insufficient expression of PEX11β. Indeed, treatment of HD cells with 4-PBA significantly improved cell viability compared to untreated controls across multiple time points (Figure 4C).

These findings suggest PEX11β is the converging point regulating co-expression of Complex II as well as MFN1/2 in HD, linking mitochondrial respiratory function to mitochondrial morphology. Balanced expression of PEX11β is also closely associated with cell survival in HD. To our knowledge, this is the first experimental demonstration of an important co-expression link between PEX11β and complex II in a pathological context such as HD. Additionally, these findings also highlight the development of potential therapeutics towards enhancing PEX11β expression to mitigate mitochondrial dysfunction with subsequent reduction of cell death in HD pathology.

### 2.5. PEX11β expression rescues the behavioral defect of 3-NP HD mice model with subsequent improvement of complex - II as well as mitochondrial fusion gene expression

Previous biochemical analyses on post-mortem HD brain tissue, along with studies in model organisms, have consistently revealed a significant and selective impairment in mitochondrial complex II activity. Notably, this defect appears to be specific, as complex I (NADH dehydrogenase) activity remains unaffected in HD brain samples(*44*) (*45*). In the chemical model of HD, experimental administration of 3-NP (3-nitropropionic acid), a selective inhibitor of COMPLEX II, has been shown to preferentially damage striatal neurons in non-human primates exhibiting the pattern of neurodegeneration of HD patients as well as behavioral defects like motor dysfunction and cognitive impairments (*46*) (*47*). This neurochemical environment contributes to mitochondrial dysfunction, which in turn leads to cognitive decline, motor impairments, and behavioral abnormalities commonly observed in patients with HD.

These observations prompted us to examine the expression pattern of complex II following systematic administration of 3-NP using our previously published protocol (*48*).

Consistent with earlier reports, 3-NP administration to C57BL/6 mice (n = 5) resulted in pronounced motor dysfunction (Figure 5A, D). Cognitive impairment was also evident, as demonstrated by poor memory performance in the water maze test (Figure 5A, D See video supplementary file). We assessed complex II expression pattern in the striatum–the primary site of neurodegeneration in HD—and the hippocampus, a region critical for learning and memory. Both regions exhibited reduced expression of complex II (SDHB), indicating mitochondrial dysfunction (Figure 5B, Supplementary figure 5A). Notably, similar to our *in vitro* genetic model, we observed a significant reduction in PEX11β expression in the striatum and hippocampus, suggesting a shared regulatory axis between PEX11β and OXPHOS (complex II) gene expression in HD pathophysiology (Figure 5B, Supplementary figure 5A). Interestingly, expression patterns of mitochondrial dynamics genes–those involved in fusion and fission–observed in the HD mouse model recapitulated the findings from our *in vitro* cellular HD model data (Figure 5C). This consistency suggests that disrupted mitochondrial dynamics are linked with altered co-expression between OXPHOS (complex II) components and the PEX11β gene.

**Figure 5:**
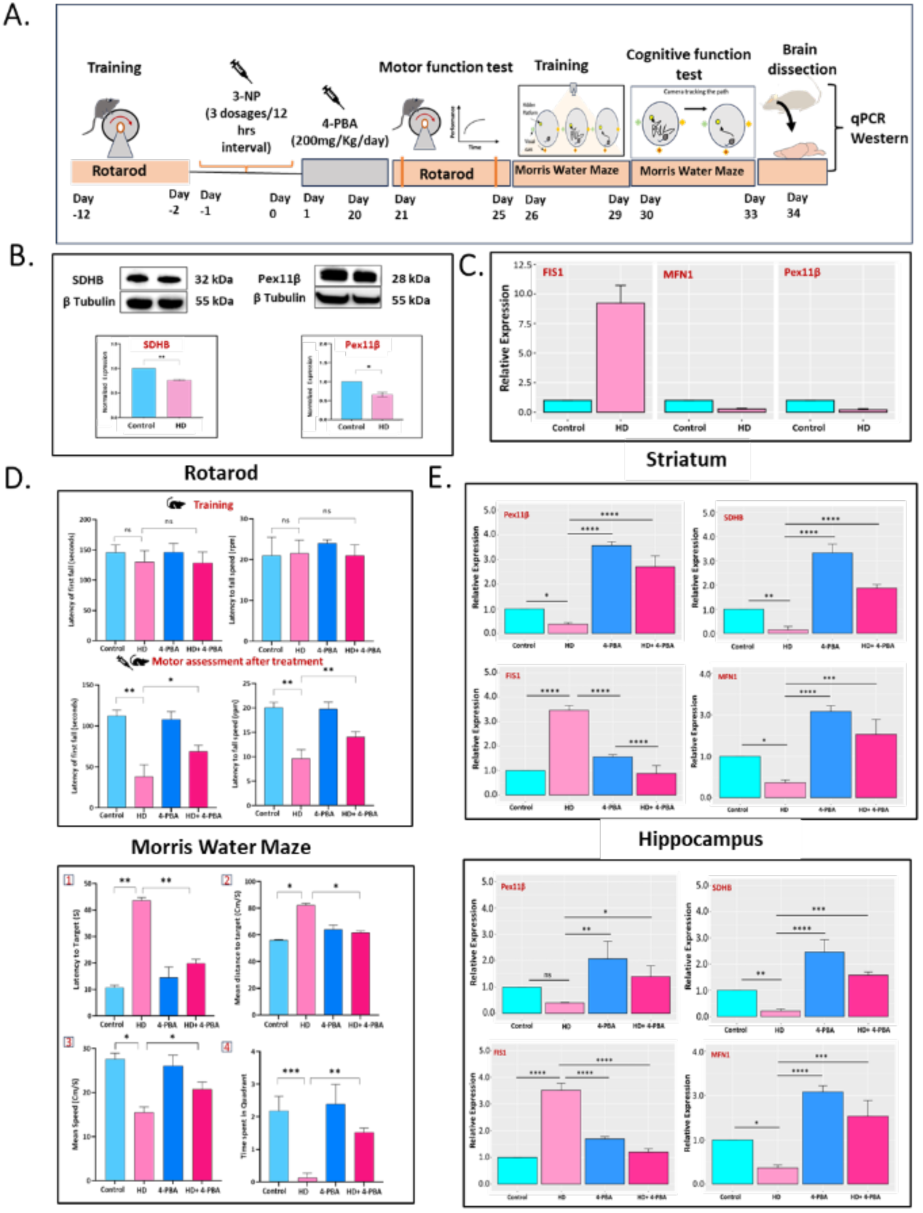
4-PBA supplementation restores expression of Pexlip and complex II in striatum and hippocampus with improved motor and cognitive behavior. (A) Schematic representation of the 3-NP-induced HD mouse model subjected to behavioral study (rotarod and Morris water maze test) to assess the defect in locomotory and cognitive functions at the indicated time points. Brain (striatum and hippocampus region) dissection was performed for downstream qPCR and western analysis to investigate gene and protein expression patterns, respectively. (Bj Western analysis of the complex II subunit, SDHB, showed significant down-regulation in the striatum of the HD mouse brain compared to the control (n=3). (C) qPCR analysis of the striatum of 3-NP HD mice showed enhanced expression of mitochondrial fission gene (FIS1) with decreased expression of fusion gene (MFN1), along with downregulated expression of PEXllp. All protein and gene expression data represented mean (n = 3, biological replicates) with SD. * p-value calculated via paired t-test. (D) Rotarod test (Upper panel) showed motor performance during the training phase (before treatment), and motor assessment after treatment (lower panel) (n=5). Morris Water Maze shows the learning phase across successive training days (Panels 1-3) as well as probe trial evaluating memory retention (Panel 4), indicating significant improvement of motor coordination and spatial learning of 4-PBA treatment groups (n=5). (E) The gene expression pattern of Pexlip, SDHB, and MFN1 after 4-PBA treatment was enhanced in both striatum and hippocampus, corroborating our in vitro data (n=3). The fission gene (FIS1) was downregulated after 4-PBA treatment in HD mice. For all the nice data, statistical significance was represented by the mean with SD. *p value calculated via paired Student’s t-test (two-tailed); * represents the following p values: *p < 0.05. **p < 0.005, ***p < 0.0005, and ****p < 0.00005.

If PEX11β is a key regulator in maintaining the fission-fusion dynamics essential for maintaining a healthy mitochondrial network and behavioral function, then restoring its expression should alleviate the behavioral deficits observed in 3-NP-induced HD mice model. To test this, we administered 4-PBA intraperitoneally prior to 3-NP and conducted behavioral assessments as described in the methods section (Figure 5A). Remarkably, 4-PBA treatment rescued both motor dysfunction as well as short-term memory impairments associated with 3-NP-induced HD symptoms (Figure 5D, Supplementary figure 5B). Molecular analyses confirmed that 4-PBA restored PEX11β expression in the striatum and hippocampus, accompanied by improved expression of complex II subunits and mitochondrial fusion genes, indicating enhanced mitochondrial health (Figure 5E, Supplementary figure 5C). Collectively, these findings underscore the critical role of PEX11β in regulating mitochondrial remodelling and its association with the co-expression of complex II components linking behavioral phenotypes with HD pathogenesis.

### 2.6. Ying Yang 1 (YY1) is a transcriptional regulator balancing the co-expression nexus of complex II and PEX11β

The transcript level co-expression regulation of PEX11β and complex II is consistent with a recent study on the co-regulation landscape of the human proteome (*19*) (*40*). Given the significant downregulation of PEX11β observed alongside disrupted mitochondrial dynamics and a characteristic decrease in nuclear-encoded complex II genes (Figure 2B, Figure 4A, Supplementary Figure 6A), we sought to explore potential regulatory mechanisms underlying this co-expression pattern. Interestingly, unlike other respiratory complexes, subunits of complex -II are encoded only by the nuclear genome (*49*). This indicates the involvement of a common transcriptional regulator controlling nuclear DNA encoded genes of complex II and PEX11β. We first investigated whether both PEX11β and complex II genes share any common transcription factor binding motif(s) in upstream of the transcription start site. Studies in both humans and mice have demonstrated that a substantial number of genes involved in OXPHOS not only interact at a protein level but also exhibit coordinated mRNA expression across various conditions and species (*50*) (*51*) (*52*) (*53*). Given that neurons predominantly use OXPHOS systems to meet their energy demands, precise regulation of this system is critical. In mammals, a large-scale *in silico* analysis by Waveren and Moraes identified significant enrichment of binding motifs of four different transcription factors in OXPHOS genes, including YY1 (*54*). Consistent with this, our own transcription factor enrichment analysis using the X2K web platform on 209 mitochondrial differentially expressed genes (mtDEGs) identified YY1 among the top three enriched transcription factors (Figure 6A, Supplementary File IV). Notably, among the genes controlled by YY1, a significant proportion (16 out of 49 genes, i.e., 32.6%) were mitochondrial OXPHOS genes, including SDHB.

**Figure 6:**
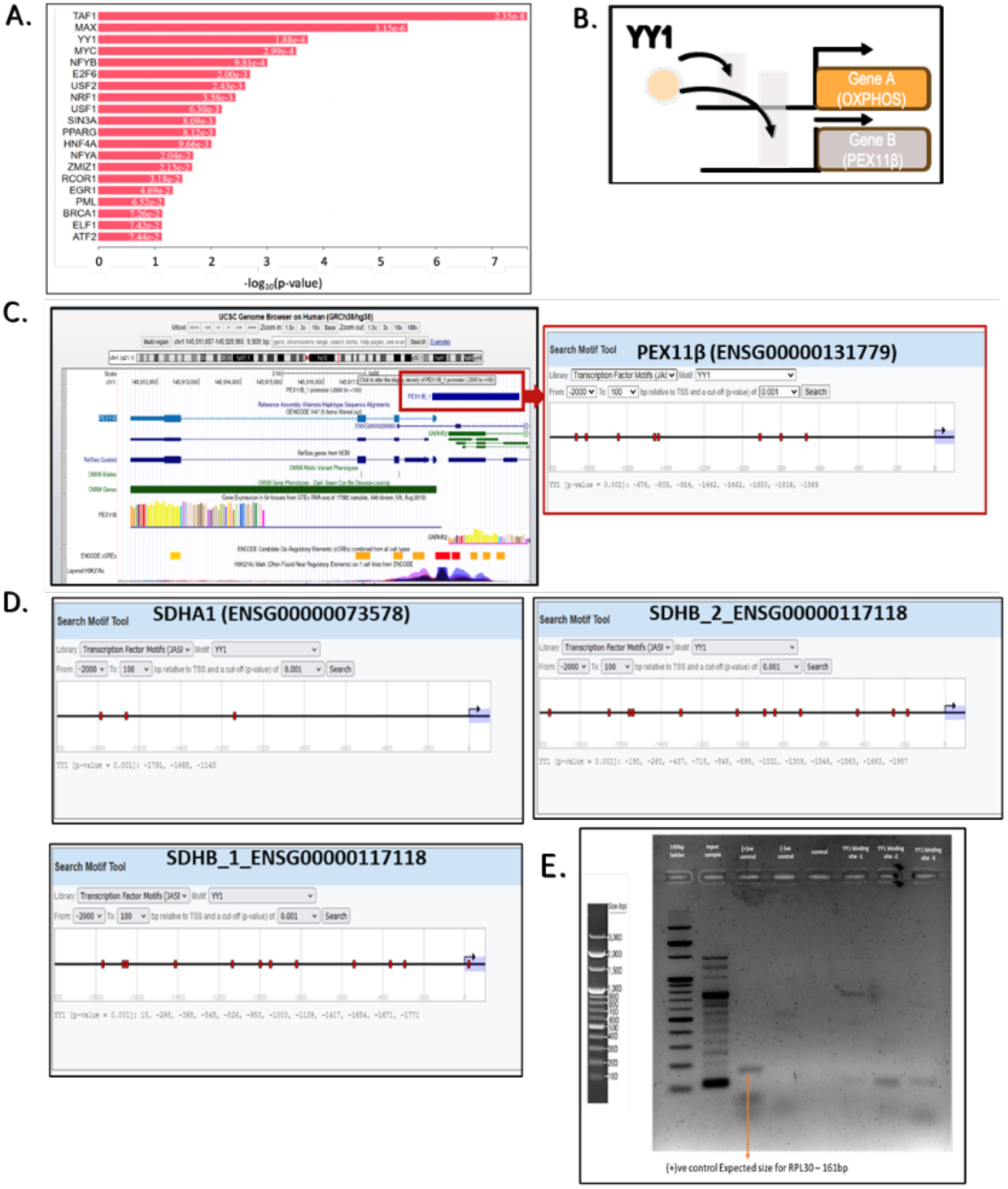
I**d**entification **of YY1 transcription factor binding of PEXllp** (A)Transcription factor analysis using the X2K webtool over 209 common genes shared between MitoCarta3.0 and AD or HD datasets identified YY1 as a key transcription factor. (B) Diagrammatic representation of the functional model of YY1 regulation over SDHB (OXPHOS gene) and PEXlip. (C) UCSC genome browser snapshot showing the promoter sequence −2000 to +100 of human PEXllp (ENSG00000131779), with corresponding EPD (Eukaryotic promoter database) search motif tool showing four YY1 transcription factor binding sites on the Pexlip promoter region. (D) EPD search motif tool showing YY1 transcription factor binding sites for Complex II genes SDHA1 (ENSG00000073578) and SDHB1/2 (ENSG00000117118). (E) ChlP-PCR analysis confirming YY1 binding to the upstream of the Pexll p gene (lane 6). All gene expression data from A and B represented the mean (n = 3, biological replicates) with SD. *p-value calculated via paired t-test with “Bonferroni” correction.

This observation aligns with previous findings identifying YY1 as a component of the transcriptional complex that directly binds to promoter regions along with the co-factor PGC1α to regulate mitochondrial gene expression (*55*). PGC1α is also known to activate the expression of mitochondrial fusion genes such as *MFN1* and *MFN2*. In this study, we demonstrated that the reduced expression of PEX11β, complex II subunits, and fusion genes (*MFN1/2*) in the HD model was effectively rescued by 4-PBA treatment (Figure 3F). These observations indicate YY1 could be the transcription factor that regulates fusion gene expression interacting with the transcription co-factor PGC1α. Taken together, these findings point to YY1 as a key transcriptional regulator orchestrating the expression of *PEX11β*, OXPHOS and mitochondrial fusion genes (Figure 6B).

We next investigated whether the promoter region of *PEX11β* contains binding motifs for the transcription factor YY1. *In silico* analysis of the −2000 to +100 bp region relative to the transcription start site (TSS) of *PEX11β* revealed four putative YY1 binding sites (Figure 6C). Consistent with the observed co-expression patterns, multiple YY1 binding motifs were also identified in the upstream regulatory regions of SDHA and SDHB, including their promoter regions. These results strongly suggest that YY1 transcriptionally regulates the coordinated expression of PEX11β and OXPHOS genes such as SDHB (Figure 6D). Chromatin immunoprecipitation followed by qPCR (ChIP-qPCR) in *in vitro* HD models confirmed YY1 binding upstream of the PEX11β gene (Figure 6E). Furthermore, a similar expression pattern was observed in the 3-NP-induced HD mouse model, reinforcing the involvement of a YY1–PEX11β–OXPHOS regulatory axis in HD pathogenesis (Supplementary Figure 6C). Collectively, our data provide the first preliminary evidence of YY1-mediated co-regulation of PEX11β and OXPHOS genes, linking this transcriptional network to mitochondrial dysfunction in HD pathology.

A close inspection of transcriptional pattern of YY1 showed a reduction in HD condition, which was enhanced after 4-PBA treatment mirroring the restoration of PEX11β-Complex II transcriptional axis. (Supplementary Figure 6B, 6C). A marked reduction in YY1 expression was observed, which closely paralleled the downregulation of OXPHOS and mitochondrial fusion genes under disease conditions. Notably, treatment with 4-PBA led to a significant upregulation of YY1 expression, mirroring the restoration of PEX11β levels reported earlier (Figure 3F). As expected, the expression of SDHB and MFN1/2 was also restored following 4-PBA treatment (Figure 3F and 4A). These findings strongly support a coordinated expression pattern among OXPHOS genes, PEX11β, and YY1, and highlight YY1’s regulatory influence over mitochondrial fusion gene expression.

### 2.7 PEX11β-mediated balance of mitochondrial dynamics conserved in *in vitro* AD model

To determine whether the regulatory relationship between PEX11β, complex II, and YY1 is conserved in AD, we extended our investigation to an AD model. *in silico* transcriptional profiling had already indicated reduced expression of PEX11β in AD (Supplementary Figure 1C). To experimentally validate this, we transiently expressed a plasmid encoding Aβ-42 to mimic AD-like conditions (Figure 7A). Consistent with our computational findings, we observed a marked decrease in PEX11β gene levels (Figure 7B), along with enhanced expression of fission genes (DRP1, MFF) with reduced expression of fusion gene (MFN1) and the SDHB subunit of complex II (Figure 7C, E).

**Figure 7:**
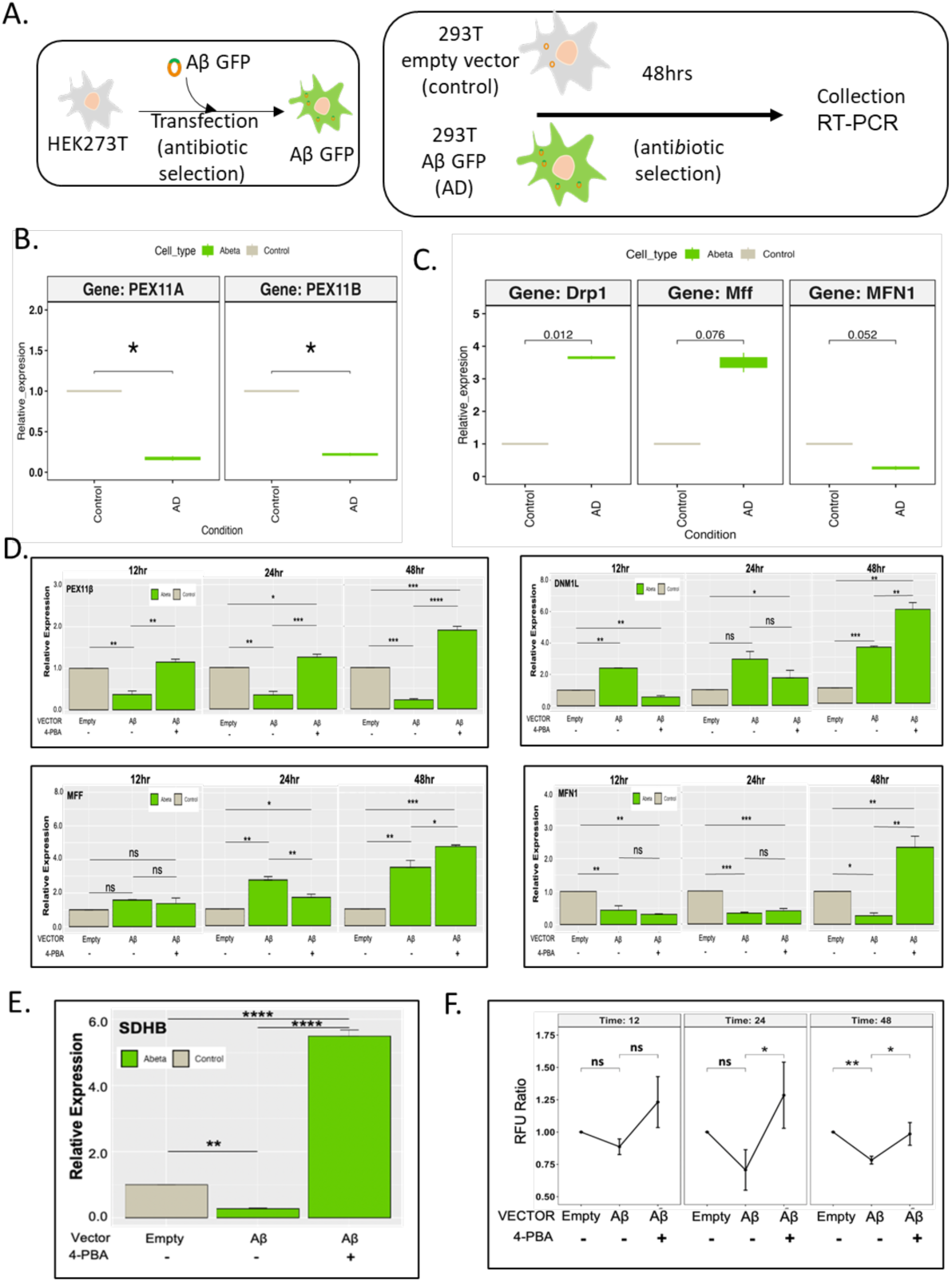
Impaired PEXlip expression and expression defect of complex II subunit and fission fusion genes in AD. (A) Schematic representation of experimental design using HEK293T line transfected with Ap-GFP for various gene expression studies. (B) RT-qPCR confirmed the downregulated expression pattern of PEXllp with transcriptomics data (n=3). (C) The expression pattern of genes controlling mitochondrial dynamics (DRP1, MFF, MFN1) is significantly downregulated in the AD sample compared to the control (n=3). (D) Bar plots across different time points of HEK293T-Ap line after 4-PBA treatment showed significant enhancement of PEXlip expression at the indicated time points (n=3). The expression pattern of genes controlling mitochondrial dynamics has also been monitored. 4-PBA seems to have significantly increased both fission and fusion genes at the later time points (n=3). (E) SDHB expression significantly increased after 4-PBA treatment (n=3). All the qPCR data (A-E) represented are from three biological replicates (n=3) with mean and SD. *p-value calculated via paired t-test. *p < 0.05, **p < 0.005, ***p < 0.0005, and ****p < 0.00005). (F) Alamar Blue cell viability assay demonstrated rescue of cellular viability in HEK293T-Ap line after 4-PBA treatment. Data collected from three independent biological replicates (n=3) were used for calculating the p-value using a paired t-test.

Expression profiling of mitochondrial dynamics genes confirmed the fusion-fission imbalance. These results align with transcriptomic analyses from HD and AD patient datasets. Treatment with 4-PBA effectively restored PEX11β expression, along with enhanced expression of SDHB and fusion genes, indicating a recovery of elongated mitochondrial morphology (Figure 7D-E). Notably, YY1 expression followed a similar pattern, as observed in the *in vitro* HD model (Supplementary Figure7A) indicating the conserved transcriptional nexus of YY1-PEX11β*-* Complex II. Collectively, these findings establish PEX11β as a critical modulator of mitochondrial fission–fusion balance, co-regulated with complex II genes via the YY1 signaling axis and extend its relevance to AD pathophysiology.

## 3. Discussion

Several recent studies have revealed a crosstalk between mitochondria and peroxisomes, highlighting their coordinated clustering and structural dynamics. However, research on the dysregulation of these cross-compartmental communications and its pathological consequences remains limited. Using a combination of *in silico* and experimental approaches, we for the first time identified how the expression defect of the peroxisomal protein PEX11β directly affects mitochondrial dynamics in two neuropathies (HD and AD) characterized by the accumulation of cytoplasmic aggregations—a hallmark of their pathological phenotype. Moreover, we demonstrated that the expression of PEX11β correlates with the expression of respiratory complex II. Restoration of PEX11β expression can cure the behavioral defect of 3-NP (inhibitor of complex II) induced HD mice, linking the molecular-level interrelation between PEX11β and complex II. Further, our extended investigation uncovers a unique transcriptional regulation mediated by the YY1 transcription factor. Overall, the presence of PEX11β is essential for cellular survivability in both disease conditions (Figure 8).

**Figure 8.**
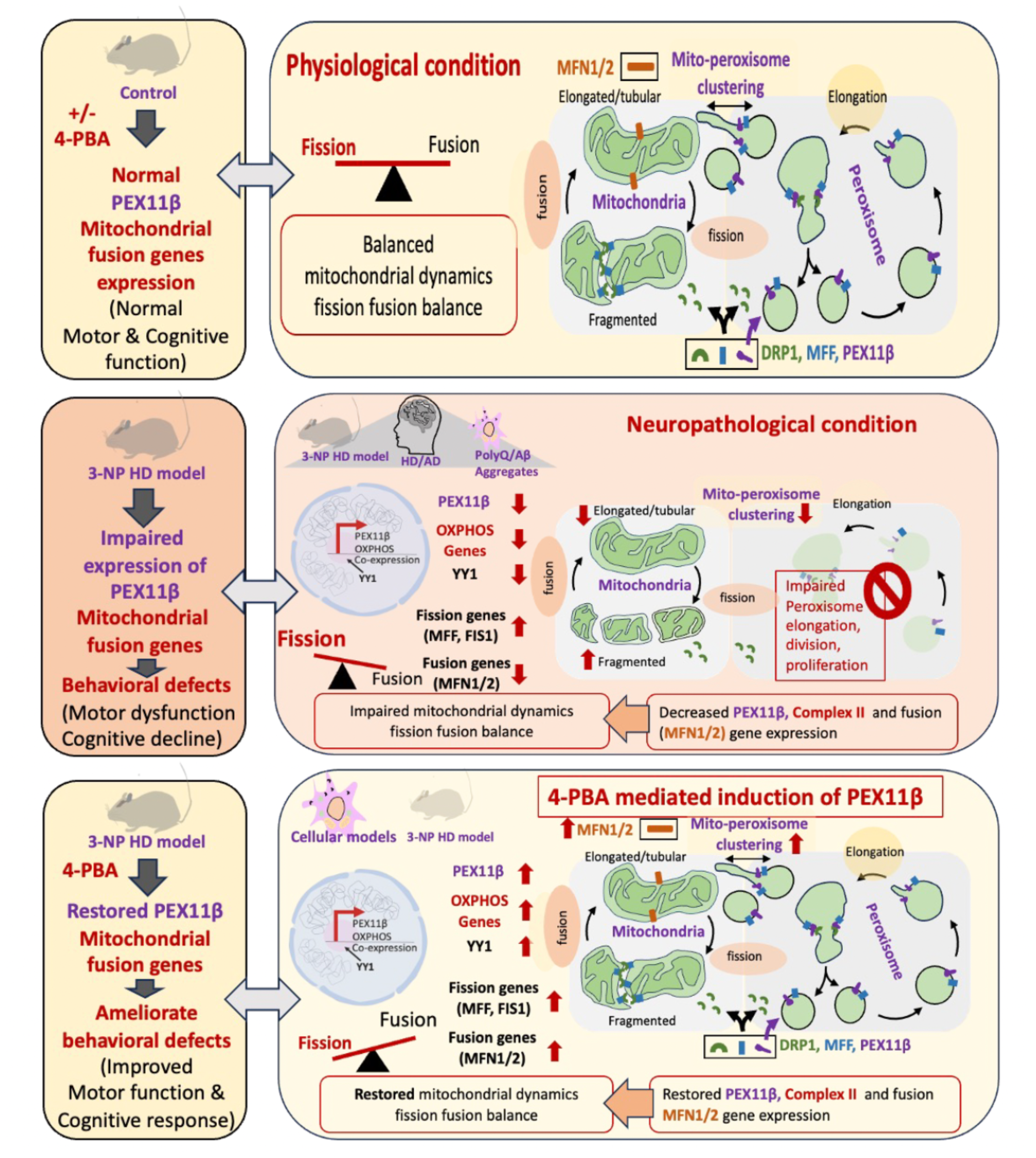
Proposed model for PEXlip mediated control over cellular phenotype (mitochondrial fusion defect) to behabioral defect during HD conditions. Schematic summarizing the molecular interrelation between PEX110 and mitochondrial dynamics in neurodegenerative conditions. Decreased expression of PEX110 directly linked to mitochondrial dynamics and complex II expression observed in Huntington’s (HD) and Alzheimer’s (AD) cellular models. Behavioural defect in 3-NP-induced HD mice model is also linked to downregulation of PEX110 and mitochondrial fusion genes. Restoration of PEX110 expression rescues expression of mitochondrial dynamics and ameliorates behavioural deficits in 3-NP-induced HD mice. The study further identifies a transcriptional link between YY1-PEX110-Complex II expression axis crucial for mitochondrial health and cellular survival in these disease condition.

Our interest in PEX11β and its relation to neuropathological conditions like HD/AD first arose when our transcriptomic analysis showed that PEX11β is one of the unique genes identified under organellar fission-fusion processes (Organelle fission GO:0048285; Mitochondrial Fission GO:0000266; Peroxisome Fission GO:0016559) (Figure 1D, Supplementary Figure 1B). In our *in silico* and *in vitro* analyses, we also observed that, along with decreased expression of PEX11β, there is a significant decrease in the expression of Complex II and other OXPHOS genes (Supplementary Figure 1C, Figure 2B, Figure 4A, Supplementary Figure 4B, C Figure 7B, Figure 7E). Though decreased Complex II expression has already been shown by several earlier studies on HD/AD, including recent cell type-specific transcriptomics analyses by Lee H et al. (*23*)(*56*) (*57*) (*10*) (*58*).

When we performed confocal imaging in *in vitro* model of HD, we observed severely fragmented mitochondria (Figure 2D). It is well known that mitochondrial fragmentation is controlled by a set of fission proteins (FIS1, MFF, DRP1). Interestingly, these mitochondrial fission fusion proteins along with PEX11β also control the peroxisomal proliferation cycle (*59*). In mammals, PEX11β plays a crucial role in promoting peroxisome elongation and division (*60*) (*61*) (*62*). Importantly, to initiate peroxisomal fission, the peroxisome must become elongated facilitated by PEX11β, followed by peroxisomal fission aided by two fission proteins, FIS1 and MFF. These proteins localize to peroxisomes to recruit DRP1/DNM1L from the cytoplasm (Itoyama et al, 2013) (*16*) (*64*).

Notably, live-cell imaging studies have demonstrated that elongated peroxisomes, compared to their spherical counterparts, engage more frequently in stable interactions with mitochondria—highlighting the importance of PEX11β in facilitating peroxisomal elongation and protrusion formation (*19*). Beyond PEX11β, recent research has shown that mitochondrial fusion proteins (MFN1/ MFN2) are essential for establishing peroxisome-mitochondrial coordination. For instance, the fusion protein Fzo1 (a homolog of mammalian mitofusins in budding yeast) naturally localizes to peroxisomes and promotes interaction with mitochondria (*65*). Enhanced expression of MFN1 and MFN2, mammalian mitochondrial fusion proteins, augments mitochondrial clustering and co-clustering of mitochondria and peroxisomes (Huo et al, 2022). Together, PEX11β, along with fusion proteins, maintains mitochondrial-peroxisomal contacts. Beyond sharing common machinery for their division, peroxisomes coordinate with mitochondria in key metabolic processes such as β-oxidation, phospholipid biosynthesis, and ROS metabolism (*67*).

To understand the interrelationships of PEX11β and mitochondrial fission-fusion proteins in HD and AD, we monitored their gene expression patterns across different time points using in vitro models (Figure 3F, Figure 7C, D). Confocal images of enhanced fragmentation of mitochondria corroborate higher expression of fission proteins (DRP1/DNM1L, MFF) along with the downregulation of mitochondrial fusion protein MFN1 in both neuropathic conditions. Further, low expression of PEX11β indicates potential dysregulation at mitochondrial-peroxisomal contacts as well as decreased peroxisomal proliferation with enhanced mitochondrial fragmentation.

To mimic disease-associated PEX11β deficiency, we performed siRNA-mediated knockdown of PEX11β in HD control cells (Q23). This intervention recapitulated the HD phenotype, and mitochondrial fragmentation, confirming the direct role of PEX11β in maintaining mitochondrial integrity (Figure 3A-C, Supplementary Figure 3A, B).

Conversely, we enhanced PEX11β expression in HD cells using 4-phenylbutyric acid (4-PBA), a chemical chaperone known to stimulate peroxisomal proliferation. Over 48 hours, we tracked the expression of PEX11β, DRP1/DNM1L, FIS1, MFF, and MFN1/2 in the presence and absence of 4-PBA and identified a unique pattern. Adding a peroxisome proliferator enhanced the expression of all genes, indicating restoration of peroxisomal division and proliferation in HD cells. Interestingly, in addition to enhanced expression of the fission proteins, we observed upregulation of the mitochondrial fusion protein MFN1/2. In agreement with this observation, our confocal imaging also showed more elongated mitochondria after adding 4-PBA in HD cells, suggesting a shift towards enhanced mitochondrial fusion and potential improvement of peroxisome-mitochondrial contacts (Figure 3E-F, Supplementary Figure 3C-E, Figure 7D).

Previous studies in mice lacking the PEX11β gene exhibited severe neurological dysfunction, defects in neuronal migration, and developmental delay (*68*). Patients with homozygous mutations resulting in non-functional truncated PEX11β have been reported to exhibit intellectual disability (*69*). All these reports underscore the critical role of PEX11β as a key regulator of peroxisome maturation and division, with its dysfunction linked to neuropathological conditions.

In HD, apart from disrupted fission-fusion dynamics, mitochondrial dysfunction has been linked to defects in mitochondrial protein import due to the direct interaction of mutant huntingtin with the mitochondria, impaired biogenesis, excessive ROS production, and mtRNA-mediated downregulation of oxidative phosphorylation (OXPHOS) genes (*70*). In AD, amyloid aggregates have been shown to have similar alterations, such as altered membrane potential, ROS production, and dysfunction of OXPHOS (*71*) (*72*). Due to limited glycolytic capacity, neurons rely heavily on mitochondrial oxidative phosphorylation (OXPHOS) for energy production (*1*) (*2*) (*3*). Our differential gene expression (DEG) analysis also captured this phenomenon, where we observed significant enrichment of ROS-related pathways in both diseases (GO:0000302, GO:2000379, GO:0032928, GO:0032930) (Supplementary Figure 1C). Experimental validation confirmed elevated ROS levels and altered membrane potential in HD and AD cell models (Supplementary Figure 2B, C). Impaired OXPHOS and increased ROS production contribute to disease pathology.

It is important to note that Complex II is unique among respiratory complexes. Complex II comprises nuclear-genome encoded subunits and does not pump protons across the mitochondrial membrane (*49*) (*73*) . It transfers electrons from succinate to ubiquinone and plays a role in redox regulation. Impairment in Complex II disrupts electron flow to Complex III, leading to oxidative stress. Dysfunction of Complex II has been identified as a key pathological feature in both HD and AD (*44*) (*45*). In HD, postmortem brain samples show reduced Complex II expression, and both *in vitro* and *in vivo* studies link striatal neurodegeneration to Complex II defects(*4*).

We aimed to explore the link between Complex II and PEX11β to understand organelle crosstalk in HD and AD. A recent human proteome co-regulation map showed that PEX11β exhibited co-expression with ATP synthase and other ETC components (*20*) (*51*). Further, Kustatscher et al. (2019) demonstrated that PEX11β is involved in peroxisome-mitochondria interactions in COS-7 cells (*19*). Live-cell imaging revealed that elongated peroxisomes—dependent on PEX11β—form more stable contacts with mitochondria via Miro1 (*19*).

Not only PEX11β, but recent studies have shown that mitochondrial fusion proteins are required for peroxisome-mitochondrial interaction. For example, in budding yeast, the fusion protein Fzo1 (yeast mitofusin, mammalian homolog) naturally localizes to peroxisomes and promotes peroxisome-mitochondrial contacts (Alsayyah et al., 2024). Enhanced expression of MFN1 and MFN2, mammalian mitochondrial fusion proteins, augments mitochondrial clustering as well as co-clustering of mitochondria and peroxisomes (Huo et al, 2022). Importantly, peroxisomal β-oxidation and mitochondrial ATP generation are found to be linked, and peroxisome–mitochondria contact sites are essential to maintain this process (*74*). In line with these observations, we showed a co-expression similarity between PEX11β and complex II in the HD/AD condition (Figure 2B, Figure 4A, Supplementary Figure 4B). The treatment of the peroxisome proliferator, 4-PBA, enhanced PEX11β expression, which rescued the decreased gene expression coding for complex II subunits (Figure 3F, Figure 4A). Importantly, 4-PBA treatment cannot restore complex II subunit expression in the siRNA-mediated PEX11β-knockdown condition (Figure 4B). Thus, the presence of PEX11β is essential in maintaining complex II expression. If 4-PBA restores energy metabolism by improving complex II expression, it should help cellular survivability. We confirmed enhanced survivability in the presence of 4-PBA in both HD and AD conditions (Figure 4C, Supplementary Figure 7F). To our knowledge, we are demonstrating a novel co-expression axis between PEX11β and Complex II for the first time, revealing a new layer of signaling critical to HD/AD pathogenesis. Moreover, the FDA-approved prodrug 4-PBA enhances survivability by restoring PEX11β expression and mitochondrial dynamics, offering a promising therapeutic strategy.

Earlier studies with mice lacking the PEX11β gene exhibited severe neurological dysfunction, neuronal migration defects, and developmental delay (*68*). It is interesting to note that patients harboring the PEX11β mutation (c.743_744delTCinsA mutation in the exon 4 of the PEX11 β gene), causing a subgroup of peroxisomal biogenesis disorders, exhibited symptoms of neuronal anomalies with abnormal gait (*17*). Strikingly, the treatment regime targeting to improve mitochondrial function ameliorated the symptoms, suggesting the involvement of a complementary functional relationship between PEX11β and mitochondrial dysfunction (*17*). However, the participation of PEX11β in causing mitochondrial dysfunction in neurodegenerative disease is largely unknown. Abnormal gait and mitochondrial dysfunction are also hallmark symptoms of HD (*48*) (*75*). To assess the conservation of PEX11β and mitochondrial dysfunction (Complex II expression defect, ROS production, etc) signaling axis and its behavioral impact, we conducted *in vivo* studies using a targeted Complex II inhibition model of HD (Figure 5A). Previous research showed that 3-nitropropionic acid (3-NP) induced chemical model of HD exhibits mitochondrial dysfunction and ROS production, causing striatal neuron degeneration. As observed in HD patients, 3-NP-induced HD mouse exhibits motor deficits and abnormal gait (*48*).

In this model, we confirmed that reduced Complex II expression correlates with downregulation of PEX11β and mitochondrial fusion genes in the striatum (Figure 5C, E, Supplementary figure 5C) at the molecular level. If PEX11β is the master regulator of the PEX11β -complex II co-expression axis and is linked to behavioral symptoms, restoring PEX11β will improve behavioral defects. Indeed, 4-PBA treatment to 3-NP HD mouse model restored PEX11β expression as well as SDHA/SDHB expression (Complex II subunits) with subsequent cure of motor and cognitive deficits (Figure 5D, Supplementary figure 5B, see video supplementary file).

Finally, we investigated the transcriptional regulation of this co-expression axis. No prior studies have defined how PEX11β and Complex II are co-regulated. To this end, through *in silico* analysis, we focused on a set of potential transcription factors known to control neurogenesis and maintain homeostasis in the developing brain. Using X2K web-based transcription factor enrichment analysis on 209 mtDEGs, we identified YY1 among the top regulators. A previous ChIP-seq study demonstrated that mitochondrial genes are under the control of YY1 (*55*). Similarly, earlier reports of in silico analysis revealed a positive correlation between YY1 and OXPHOS gene expression (*54*). In addition, knockdown of YY1 showed defective mitochondrial morphology and bioenergetic function in mouse skeletal muscle (*76*). Our *in silico* transcription factor enrichment analysis with X2K web using the 209 mtDEGs identified YY1 among the top 3 transcription factors (Figure 6A). In addition, we mapped potential binding sites of YY1 at the upstream of PEX11β and complex II genes (Figure 6B-D). Earlier reports on HD/AD pointed out PGC-1α as a transcriptional coactivator involved in regulating transcription of mitochondrial gene expression (*76*) (*77*) (*78*). Interestingly, PGC-1α acts as a transcriptional coactivator in conjunction with different transcription factors, including YY1 (*78*). Thus, YY1 with PGC-1α can control the expression of mitochondrial genes in HD/AD conditions. Indeed, our ChIP-qPCR data confirmed the binding of YY1 at the promoter region of PEX11β as well as complex II genes (Figure 6E). To our knowledge, identifying YY1 binding to PEX11β is also the first report. These observations established a potential transcriptional co-expression governed by YY1 and improved our understanding of how transcriptional control can influence peroxisome-mitochondrial communication during HD or AD pathology.

In conclusion, our integrative analysis reveals that PEX11β plays a central role in peroxisome-mitochondrial crosstalk and is a key regulator of mitochondrial dynamics. We identified a novel co-expression axis linking PEX11β and Complex II, governed by the transcription factor YY1. These findings highlight the multifaceted role of PEX11β in maintaining mitochondrial health and its significance in HD and AD pathology. The therapeutic potential of 4-PBA in restoring PEX11β expression and mitochondrial function opens new avenues for treating neurodegenerative diseases.

## 4. Materials and methods

### 4.1. Data collection

We used two datasets for HD (GSE33000, GSE64810) and three datasets for AD (GSE125583, GSE132903, GSE33000) based on the following criteria: Publicly available data from the previous 10 years of study, b. Omics data from brain tissues only, c. A minimum sample size of 50 (for HD)/ 150 (for AD). Since there was only one HD dataset (GSE33000) available with more than >150 samples, we have lowered our minimum sample size criteria to 50, d. Animal model studies were excluded, and e. Compatibility with GEO2R-based analysis. Detailed descriptions of the datasets are provided in Supplemental Table I.

### 4.2 Data analysis

We employed the GEO2R (*79*) tool to detect differentially expressed genes (DEGs). GEO2R log-transformed the data, and then p-values were adjusted based on the Benjamini & Hochberg (False Discovery rate, FDR) method for multiple hypothesis testing. For AD, 3293 genes were common (considering genes shared by at least two datasets) from the top 5000 genes (2,500 upregulated and 2,500 downregulated with a P-adjusted value of <0.05) identified in the three datasets (Supplementary Table II, Supplementary File II). This includes 1,497 upregulated and 1,659 downregulated genes (Supplementary Table II). Similarly, for HD, 1,082 genes (673 upregulated and 262 downregulated genes) were shared between the two datasets (Supplementary Table I & II). We identified 209 mitochondria-specific genes by intersecting genes from the MitoCarta3.0 database (1,136 genes, Supplementary File III) and common DEGs mentioned above from AD (3,293 genes) and HD (1,082 genes) datasets. Enrichment analysis was performed using Enrichr (*80*). Enrichr groups genes into different pathways using databases such as KEGG, Reactome, panther, and Gene Ontology databases for biological processes and molecular function and ranks pathways based on combined scores (calculated by multiplying the Z-score by the log of p-value derived from Fisher’s exact test).

### 4.3 Visualization and analysis of the PPI network

Cytoscape (3.10.2) was used to visualize and analyze the PPI network. We used the ClueGO plugin with Gene Ontology (GO) terms - Cellular components (*81*). This enabled us to visualize the interaction between cellular components involving DEGs identified in the upstream analysis. Through this interaction network, we identified the genes/proteins involved in particular cellular processes. Moreover, it allows us to visualize the interacting organelles involved in a particular disease condition. ClueGO employs Fisher’s Exact Test to calculate p-values, determining the significance of term enrichments, and the kappa score is utilized to define term-term relationships (edges) and functional groups based on shared genes between terms, with a kappa score threshold of > 0.4.

### 4.4. 3-NP induced Mouse HD model set up and Rotarod test

Male C57BL/6 mice, aged 10 to 12 weeks and weighing an average of 26 ± 3 g, were obtained from ICMR - National Animal Resource Facility for Biomedical Research (ICMR-NARFBR, Hyderabad, India). The animals were housed (3 mice per cage) with unrestricted access to food and water under a 12-hour light/dark cycle at a temperature of 25 ± 2°C. All animal experiments were conducted with the approval of the Institutional Animal Ethics Committee (IAEC) at BITS Pilani, Hyderabad, India (BITS-Hyd-IAEC-2022-12). The animals were randomly assigned to two experimental groups (control and treatment). The treatment group (n = 5) was administered 3-nitro propionic acid (3-NP; Cat No.: N22908, Sigma) intraperitoneally at a dose of 25 mg/kg body weight spaced 12 hours apart for a total cumulative dose of 75 mg/kg body weight. The control group (n = 5) received 0.9% saline (*48*). To test the effect of Pex11β overexpression over behavioural defect, we supplemented the mice with 200mg/kg/day of 4-PBA (Sigma Aldrich - P21005) for 20 days intraperitoneally (Figure 5A). After the 4-PBA treatment, we assessed motor coordination using the rotarod, following the method described by Kumar et al. (2009) (*82*) among all four groups (Control, HD, 4-PBA, HD+4-PBA). Using the rotarod, the latency to fall was tracked for 180 seconds and analysed to understand the differences in motor function between control and 3-NP treated mice. Along with the motor dysfunction test (Rotarod), we tested cognitive response through Morris water maze test following the established protocols described by Charles V. Vorhees and Michael T. Williams (2006) and Gallagher et al. (1993) (*83*) (*84*). Briefly, after sequential training for three days, we assessed escape latency, path length, and swimming speed of all the animals along with a probe trial (24 hours later) to evaluate spatial memory retention by measuring the time spent in the target quadrant (Figure 5D)

### 4.5 Cloning Aâ (1–42) - Green Fluorescent Protein (GFP) in expression vector pcDNA3.1 (+) for AD model

To generate the plasmid pcDNA3.1-Aβ (1–42)-GFP, the Aβ (1–42)-GFP sequence from the Aβ-GFP plasmid (Addgene Plasmid #72203) (a kind gift from Venkatesh Sivaramakrishnan) was amplified by polymerase chain reaction (PCR) using primers Aβ(1–42)-GFP Forward primer_HindIII: 5’-ttAAGCTTatggatgcggaatttcgcca-3’ and Aβ(1–42)-GFP Reverse primer_XbaI: 5’aaTCTAGAtcagtggtggtggtggtggt 3’. The PCR was carried out using the Q5 High-Fidelity DNA polymerase Enzyme Mix (NEB-M0491S) as per the supplier’s instructions. The resulting 910-basepairs (bp) PCR fragment was purified with the GenepHlow Gel/PCR Kit (DFH100-Geneaid), double digested with BamHI (R3136S-NEB) and XbaI (R0145S-NEB) and cloned into the pcDNA3.1 (+) plasmid (Supplementary file I). The insert was confirmed via PCR as well as Sanger sequencing (Supplementary file I). Plasmid preparation was performed using the HiPurA plasmid isolation kit (HiMedia Laboratories Private Limited, India) for transfection into the HEK293T cell line.

### 4.6 Cell lines, culture and experimental set-up for *in vitro* HD and AD models

Recombinant HEK293 cell lines (containing exon 1 of the human HTT gene, with either 23 CAG (control; Q23) or 74 CAG (HD; Q74) repeats, fused to GFP under the control of a doxycycline-inducible promoter) were generous gifts from David C Rubinsztein’s lab, University of Cambridge, UK (*85*). Both cell lines were negative for mycoplasma contamination as tested by the EZdetect PCR Kit (HiMedia Laboratories Private Limited) (Supplementary file I).

The Q23 and Q74 cell lines were cultured in poly-l-ornithine-coated plates in Dulbecco’s Modified Eagle’s Medium (DMEM, Gibco, Thermo Fisher Scientific, USA) supplemented with 10% heat-inactivated fetal bovine serum (Gibco, Thermo Fisher Scientific) with a cocktail of antibiotics [10000 units/ml penicillin, 10 mg/ml streptomycin (Gibco, Thermo Fisher Scientific), Hygromycin (150 µg/ml), blasticidin (5 µg/ml)] and maintained at 37°C with a 5% CO2 atmosphere. At 60% – 70% confluence, doxycycline (1 µg/ml) was added to induce PolyQ HTT formation for 48 hours. For the knockdown experiment, siRNA and scrambled were transfected as prescribed in jetPRIME® transfection protocol (PolyPlus).

For the AD study, HEK293T cells were grown in DMEM (DMEM, Gibco, Thermo Fisher Scientific) with 10% FBS and an Antibiotic mix (10000 units/ml penicillin and 10 mg/ml streptomycin) at 37°C with 5% CO2 atmosphere. At 70% confluence the cells were transfected with pcDNA3.1-Aβ (1–42)-GFP plasmid using jetPRIME® transfection kit (Polyplus) following the manufacturer’s instructions. After 12hrs the old media was replaced by antibiotic selection (Geneticin, Gibco) media.

We also utilized the pARiS system for transiently expressing Q23/Q100 repeats in Neuro2a (N2A) cells. The pARiS plasmids were a generous gift from the Frederic Saudou lab. Jet-Prime kit-based transfection (Cat.No: 101000027/0.1ml) was performed to express PolyQ plasmids as per the previously described protocol (*86*). Briefly, Neuro2a (N2A) cells were cultured in standard cell culture conditions. At 70% confluency, cells were transiently transfected with the respective plasmids and incubated for four hours, before being replaced with new media. To test the effect of Pex11β overexpression on mitochondrial morphology, and cellular viability, we treated the lines with 4-PBA (5Mm, Sigma Aldrich - P21005) for 48 hours as represented in the schematic (Figure 3E).

### 4.7 Survival assay

Cell survival was determined by alamarBlue® (BIO-RAD) as per the manufacturer’s instructions. To assay the survival of the HTT and AD cell lines, 10,000 cells were seeded in each well of the 96-well plates and incubated at 37°C with a 5% CO_2_ atmosphere over a poly-L-ornithine (Sigma Aldrich) coated surface. At 60% confluency, doxycycline was added to induce polyQ HTT formation. Six hours after doxycycline treatment, 4-Phenyl butyric acid (4-PBA) was added at a concentration of 1mM and incubated for 12, 24, 48 hours. For checking the survivability, 20 µL of alamarBlue® (10%) was added to each well and incubated for 4 hours before taking fluorescence measurement. Fluorescence output was measured at the excitation wavelength of 560 nm and emission wavelength of 590 nm in a microplate reader (SpectraMax iD3, Molecular Devices). We performed three independent experiments with 3 wells for each condition. The percentage difference in reduction between treated (4-PBA) and control cells was determined. For statistical analysis, all measurements were normalized relative to the control (e.g., no treatment). For AD cells after transfection of pcDNA3.1-Aβ (1–42)-GFP plasmid, measurement of Alamar Blue fluorescence was taken after 4-PBA treatment for the indicated hours (12, 24, and 48 hours) (Figure 7F).

### 4.8. Quantitative RT-PCR

Total RNA was isolated from HTT and AD cell lines using the Trizol method (RNAiso Plus-Takara) at the indicated hours (12, 24, 48hrs), while the quantity and quality of RNA were determined using the NanoDrop (Thermo Scientific). Using the PrimeScript 1st strand cDNA synthesis kit (Takara), 1µg of total RNA was reverse transcribed to prepare the respective cDNA. RT-qPCR was carried out in 96-well plates either in the Light Cycler 480 (Roche) or in CFX96-real-time PCR machine (Bio-Rad) using the SYBR green master mix (TB Green, Takara) with specific primers. The conditions of the PCR cycle were, 95°C for 10 min, followed by 45 cycles of 95°C for 15s, 60°C for 10s, and 72°C for 10s. Three technical replicates for each gene were used for three independent experiments. Relative gene expression was quantified using the ΔCt method for all the primers using β-actin as the normalizing gene.

We used the ΔΔCt method to determine the fold changes between treatment and control groups where fold change = 2^(-ΔΔCt). Here, ΔCt (cycle difference) = Ct (target gene) - Ct (control gene), and ΔΔCt = ΔCt (treated condition) - ΔCt (control condition) (*87*). Primer details are given in Supplementary Table V.

### 4.9. Confocal microscopy

Cells were seeded in confocal dishes at a density of 1.8 x 10^6^ cells at 37°C with a 5% CO_2_ atmosphere. After 60% – 70% confluency, cells were induced with 1µg/ml doxycycline (Sigma-Aldrich) for 6 hours, followed by 4-PBA (1mM, Sigma-Aldrich) treatment for 48 hours before imaging. Before imaging, confocal dishes were transferred to the temperature and CO_2_-controlled chamber attached to the Leica confocal (SP8/DMi8) microscope. Cells were incubated with MitoTracker red (100nM, Invitrogen, Thermo Scientific) for 5 minutes at 37°C before capturing the image with the appropriate excitation/emission setting (581nm/644nm). Appropriate excitation/emission (475nm/ 509nm) was used to capture the GFP fluorescence. Fluorescence as well as corresponding bright field images were taken for multiple fields with the same setting. The images were further processed in Fiji (Image J, NIH), a freely available Java-based image processing program, to enhance the brightness and contrast (*88*). We used basic LUT adjustments, and the lookup table was made using the command: Image ------- Adjust Brightness/Contrast. The first two scrollbars (Minimum and Maximum) were used to adjust thresholds below and above, for which pixels are given the first and last LUT color, respectively. The brightness and contrast were automatically adjusted through these sliders. Different features of mitochondria were analyzed using MiNA as per the developer’s (StuartLab, https://github.com/stuartlab) instructions (*89*).

### 4.10. Western blotting

Both HD and AD cell lines were collected after 48 hours of Dox and 4-PBA treatment, lysed in RIPA buffer by continuous agitation for 1 minute and left on ice for 5 to 10 minutes. The lysate was centrifuged to remove cell debris and the supernatant was collected. The concentration of the protein was measured using a BCA protein assay kit (Pierce, Thermo Fisher). Total proteins (75µg) were separated by SDS-PAGE and transferred to a polyvinylidene fluoride (PVDF) membrane (Immobilon PVDF, IPVH00010, Millipore). The membrane was blocked in 8% bovine serum albumin (BSA) for one hour, followed by primary antibody incubation overnight at 4℃ with agitation. The following antibodies were used: PEX11β (A20953, 1:1000, ABclonal), DRP1 (A21968, 1:1000, ABclonal), Mitofusin-1/MFN1 (A21293, 1:1000, ABclonal), Mitofusin-2/MFN2 (D2D10, 1:1000, CST), RHOT1/MIRO1 (A22551, 1:1000, ABclonal), SDHB (A10821, 1:1000, ABclonal),

β-Tubulin (AC008, 1:1000, ABclonal), FIS1 (E3K90, 1:1000, CST). The next day, after washing the blots with Tris-buffered saline with 0.1% Tween 20 detergent (TBST), secondary antibody was added and incubated for one hour at room temperature with agitation. The antibody used was: Anti-rabbit IgG, HRP-linked antibody (1:10,000, 7074S, CST). The signal intensities of the bands were captured using the fusion pulse gel documentation system (Eppendorf, USA). ImageJ software was used to quantify the band intensities. Using Image J software, the protein expression was measured and normalized to the protein intensity of housekeeping gene. The details of all blots (3 replicates) can be accessed in Supplementary file V.

### 4.11. Transcription Factor Analysis

To find the YY1 transcription factor binding site at the upstream (-2000 to +100) of PEX11β and SDHB, the EPD (Eukaryotic promoter database) search motif tool was used. We also used X2K analysis tool (https://maayanlab.cloud/X2K/) to find the potential transcription factors binding across 209 mitochondria-specific genes (Supplementary File IV).

### 4.12. Chromatin Immunoprecipitation (ChIP)-qPCR

ChIP was performed by following the SimpleChIP ® Enzymatic Chromatin IP Kit (Cell Signaling Technology, Cat No: 9003S). Briefly, Q23/Q74 cell lines were cultured for 48 h in 10 cm dishes until ∼90% confluency. Further, cross-linking was performed using formaldehyde at a final concentration of 1% by directly adding it to the culture medium and by incubating for 10 minutes at 37 °C. Cross-linking was quenched with 0.125 M glycine for 5 min at room temperature. Cells were then washed twice with ice-cold PBS and harvested by scraping into PBS containing protease inhibitors, followed by centrifugation at 2000 × g for 5 min at 4 °C. Cell pellets were either processed immediately or stored at –80 °C. For nuclear preparation, cell pellets were incubated in Buffer A containing protease inhibitors and DTT on ice for 10 minutes, followed by centrifugation. Nuclei were washed twice with Buffer B and resuspended in 1X ChIP Buffer. Chromatin was digested with micrococcal nuclease at 37 °C for 20 min to yield fragments of ∼150–900 bp. Reactions were stopped with EDTA, and nuclei were lysed by homogenization. The lysates were clarified by centrifugation, and supernatants containing cross-linked chromatin were stored at –80 °C until use. 5–10 µg of digested chromatin was diluted in ChIP buffer and incubated overnight at 4 °C with the appropriate antibody. Protein G magnetic beads were then added and incubated for 2 hr at 4 °C with rotation. Beads were then washed three times with low-salt wash buffer and once with high-salt wash buffer. Chromatin was eluted from beads in ChIP elution buffer for 30 min at 65 °C. Cross-links were reversed by incubation at 65 °C for 2 h in the presence of NaCl and Proteinase K. DNA was then purified using spin columns according to the manufacturer’s instructions (Monarch DNA). Input and ChIP DNA were analyzed by PCR with primers specific for RPL30 as a control and target gene Pex11β.

### 4.13. Statistical analysis

Post-processing data analysis and statistical analyses were done in R version 4.3.3 (R Core Team, www.R-project.org) using specific packages. The CRAN R packages: “tidyverse” and “devtools” were used for basic data manipulation, visualization and statistical analysis. These packages allowed access to ‘rstatix’ (statistical calculations), ‘ggpubr’, ‘ggplot2’, ‘dplyr’, ‘tidyr’, ‘readr’, ‘purrr’, and ‘tibble’ libraries for data analysis and visualization. P-Value was calculated through pairwise t-test with default Bonferroni correction indicated in the respective figure legends. The p-value significance is denoted by an asterisk (*) (* represents the following p-values: *p < 0.05, **p < 0.005, ***p < 0.0005, ****p < 0.00005).

## 5. Declarations

### Ethics approval and consent to participate

Mouse Experiment performed following IAEC (BITS-Hyd-IAEC-2022-12) regulation.

### Consent for publication

Not applicable

### Availability of data and materials

The datasets analyzed during the current study are available in the NCBI/GEO (Genome expression omnibus). The used datasets are GSE33000 (https://www.ncbi.nlm.nih.gov/geo/query/acc.cgi?acc=gse3300), GSE64810 (https://www.ncbi.nlm.nih.gov/geo/query/acc.cgi?acc=GSE64810), GSE125583 (https://www.ncbi.nlm.nih.gov/geo/query/acc.cgi?acc=GSE125583), GSE132903 (https://www.ncbi.nlm.nih.gov/geo/query/acc.cgi?acc=GSE132903).

All analyzed data output are included in the supplementary files.

## Authors’ contributions

S.M. conceptualized and supervised the project. V.K. performed the majority of wet-lab experiments., P.K. (Pradeep Kodam) performed in silico analysis, clone generation and survivability assay. K.C. helped in the mouse related experiments and P.K. (Pragya Komal) performed brain dissection and helped in behavioural studies. A.S. performed *in silico* promoter analysis. S.M., V.K., and P.K. (Pradeep Kodam) wrote the original draft with help from A.S. All authors have read and agreed to the final version of the manuscript.

## Acknowledgements

We thank David C Rubinsztein’s lab, University of Cambridge, UK for providing Recombinant HEK293 cell lines (containing exon 1 of the human HTT gene, with either 23 CAG (control; Q23) or 74 CAG (HD; Q74) repeats, fused to Green Fluorescent Protein (GFP) under the control of a doxycycline-inducible promoter). We thank Dr. Victoria Barratt from David Rubinsztein Group for helping us to get the cell lines and sharing the details of growth conditions. We also thank Prof Frédéric Saudou for providing the pARiS plasmids as a generous gift. The contribution of Central Analytical Laboratory (CAL), Birla Institute of Technology and Science-Pilani (BITS-Pilani), Hyderabad Campus, is gratefully acknowledged for providing access to BD FACS Aria II, Leica confocal microscope (Leica SP8/DMi8), RT-PCR system (BIORAD CFX Opus 96). We also acknowledge the DDT-FIST (SR/FST/LS1-526/2012) facility of the Department of Biological Sciences, BITS-Pilani, Hyderabad campus for providing access to RT-PCR (LC480 Roche, Roche), imaging system for Western blot (Fusion Pulse 6, Vilber Lourmat). DBT Builder (BT/INF/22/SP42551/2021) for providing mouse facility to conduct behavioral tests as well as brain dissection.

